# Development of Gene Editing Strategies for Human **β**-Globin (HBB) Gene Mutations

**DOI:** 10.1101/2020.01.16.908319

**Authors:** Batuhan Mert Kalkan, Ezgi Yagmur Kala, Melek Yuce, Medine Karadag Alpaslan, Fatih Kocabas

## Abstract

Recent developments in gene editing technology have enabled scientists to modify DNA sequence by using engineered endonucleases. These gene editing tools are promising candidates for clinical applications, especially for treatment of inherited disorders like sickle cell disease (SCD). SCD is caused by a point mutation in human β-globin gene (HBB). Clinical strategies have demonstrated substantial success, however there is not any permanent cure for SCD available. CRISPR/Cas9 platform uses a single endonuclease and a single guide RNA (gRNA) to induce sequence-specific DNA double strand break (DSB). When this accompanies a repair template, it allows repairing the mutated gene. In this study, it was aimed to target HBB gene via CRISPR/Cas9 genome editing tool to introduce nucleotide alterations for efficient genome editing and correction of point mutations causing SCD in human cell line, by Homology Directed Repair (HDR). We have achieved to induce target specific nucleotide changes on HBB gene in the locus of mutation causing SCD. The effect of on-target activity of bone fide standard gRNA and newly developed longer gRNA were examined. It is observed that longer gRNA has higher affinity to target DNA while having the same performance for targeting and Cas9 induced DSBs. HDR mechanism was triggered by co-delivery of donor DNA repair templates in circular plasmid form. In conclusion, we have suggested methodological pipeline for efficient targeting with higher affinity to target DNA and generating desired modifications on HBB gene.

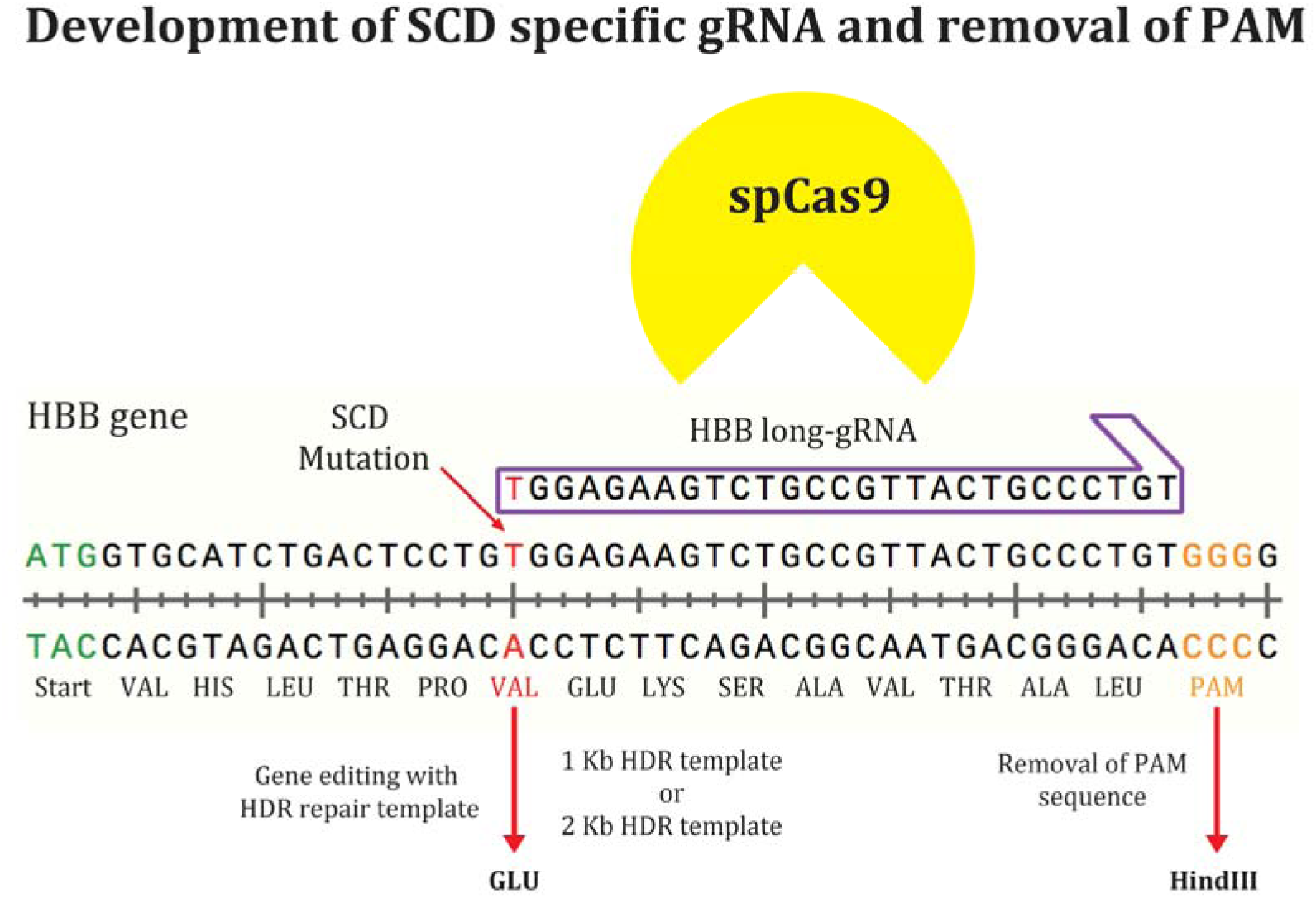
Graphical abstract

**Highlights:**

- HBB gene were targeted by spCas9 in close proximity to the SCD mutation
- Long gRNA, which is designed to target SCD mutation, is sickle cell disease specific and exhibits indistinguishable level of cleavage activity on target locus.
- Functional HBB HDR repair templates with 1 Kb and 2 Kb size were generated to cover all known mutations in the HBB gene.
- Replacement of PAM sequence in HDR template with HindIII recognition sequence allowed a quick assessment of the HDR efficiency.
- HDR template: Cas9-GFP vector 2:1 ratio yielded the highest HDR events/GFP+ cells.

## 1. Introduction

Sickle cell disease (SCD) is a devastating disease with significant morbidity and mortality rate (Gardner, 2018). The disease is characterized as a monogenic blood disease caused by a point mutation in the human β-globin gene (HBB) encoding the subunits of hemoglobin. SCD occurs due to mutated HBB allele called hemoglobin S (HbS). The point mutation (substation of A to T) in the sixth codon of HBB gene leads to the conversion of valine amino acid to glutamic acid, resulting in significant changes in the behavior and conformation of the hemoglobin molecule (Frenette and Atweh, 2007; Gardner, 2018). Abnormal HbS hemoglobin results in the collection and polymerization of the protein, which forms red blood cells in the form of sickle. Unlike doughnut shaped, elastic normal red blood cells; sickle cells have a stiff and sharp sticky structure that easily aggregate and stick on interior surface of the narrow blood vessels. As a result of occlusion, tissues are inadequately oxygenated and organ damage is seen after disease progression (Ashley-Koch et al., 2000). In addition, sickle-shaped red blood cells have a shorter life span than normal red blood cells, which may cause chronic anemia that has further pathological effects on patients health (Azar and Wong, 2017; Gardner, 2018). Clinically, the presence of homozygous variant (HbSS) is the most severe form of SCD compare to heterozygous mutations (Frenette and Atweh, 2007).

Sickle cell disease, one of the most common hereditary diseases in the world, is not only of clinical importance, but also an economic and globally vital issue (Gardner, 2018; Kapoor et al., 2018). It has been reported that around 100,000 patients have been diagnosed with SCD only in USA (Hassell, 2010; Piel, 2013; Wierenga et al., 2001). Prevalence of HgS trait and β-thalassemia trait (another common form of HBB mutation) raise up to 10% and 13.1% in some regions of Turkey, respectively (Altay and Gurgey, 1986; Beksac et al., 2011; Bircan et al., 1993; Canatan, 2014; Canatan et al., 2006; Guvenc et al., 2012; Kilinc, 2006; Mendilcioglu et al., 2011; Topal et al., 2015; Tosun et al., 2006; Zeren et al., 2007). The disease has a high prevalence especially in sub-Saharan Africa, Mediterranean basin, Middle East and India. SCD is an increasing global health problem. Estimates suggest that approximately 300,000 infants are born with sickle cell anemia each year, and this number may increase up to 400,000 by 2050 (Piel et al., 2017). Despite of high incidence of SCD, still there is not any definitive treatment for it. Current treatments are predominantly available as supportive agents to reduce disease severity and background complications. The most commonly used clinical applications in SCD patients are blood transfusion, hydroxyurea therapy and vaccinations to prevent the risk of serious infections (Aliyu et al., 2006).

Allogeneic stem cell transplantation, a promising treatment for SCD and similar blood diseases, is based on the availability of suitable donors, harvesting of healthy HSCs and their transplantation to the patients (Kahraman et al., 2014; Ozdogu et al., 2018b; Shenoy, 2011; Yesilipek, 2007; Yesilipek et al., 2018). Even though there are successful results of this method, it is not a universal treatment because of insufficiency of available donors, the immune issues like graft versus host disease (GVHD) and other toxicities (Ozdogu et al., 2018a; Ozdogu et al., 2018b). Autologous transplantation of ex vivo corrected HSCs have been proposed as a promising method to eliminate the drawbacks of allogeneic transplantation (Simsek et al., 2010; Yucel and Kocabas, 2018). Monogenic blood diseases, such as SCD, can be treated by correcting mutations directly using genome editing tools, also called engineering nucleases. These nucleases have been shown to have high potential for therapeutic applications in previous studies (Kuscu et al., 2014). Recently, type II CRISPR / Cas9 system become the most fashionable tool for genome editing and a promising approach for the direct correction of mutations that cause monogenic diseases such as SCD (Hoban et al., 2016b; Vakulskas et al., 2018).

The CRISPR/Cas9 nuclease system is composed of the DNA-binding CRISPR RNA (crRNA) array that encodes for the guide-RNA (gRNA), the auxiliary trans-activating crRNA, and the Cas9 nuclease (Hussain et al., 2019). The CrRNA contains 20-nucleotides (nt) guide sequence that binds to specific 20 base pairs DNA target in the genome by Watson-Crick base pairing. Target recognition by Cas9 nuclease depends on the protospacer contiguous motif (PAM) sequence associated with the DNA binding region.

The Cas9 nuclease induces double-stranded break three base pairs upstream of the PAM sequence. Although the CRISPR / Cas9 system is the most recently developed genome editing tool, it has recently been a striking tool for site-specific genome editing as well.

Sickle cell disease is one of the inherited diseases that is hoped to lead a cure by gene editing technology. Since it is caused by a point mutation on HBB gene, it is possible to efficiently induce base change to restore functional beta subunit of the hemoglobin. Recent studies have showed the applicability of HBB repair through application of CRISPR/Cas9 system in CD34+ HSCs (DeWitt et al., 2016). Study concluded an efficacy in gene editing for HBB locus and recovery of WT β-globin production. It still led to a detectable off-targeting activity in both HSCs and K562 cells. These studies suggested that precise and SCD specific approaches are still needed to alleviate off-targeting activity of CRISPR/Cas9 system and to upgrade HDR yield to bring gene editing technologies one step closer to clinical applications. Thus, we aimed to develop a SCD specific gRNA with high affinity to HBB locus in close proximity to SCD mutation to induce DSBs and allow repair of HBB gene with varying lengths of HDR templates to cover all possible mutations. These gene editing technologies could be also applied to other hemoglobinopathies leading to thalassemia.

## 2. Materials and Methods

### 2.1. Cell Lines and Culture Conditions

Human embryonic kidney cell line (HEK293T) was purchased from ATCC (American Type Cell Collection). HEK293T cells were cultured in high glucose (4.5 g/L) Dulbecco’s Modified Eagle Medium supplemented with 10% Fetal Bovine Serum (FBS) and 1% Penicillin-Streptomycin (Aksoz et al., 2018; Kocabas et al., 2015b). Cells were maintained at 37°C, 5% CO_2._

### 2.2. Design of gRNAs

Exact genomic location of the most common mutation (rs334) causing SCD was determined using online SNP database of National Center for Biotechnology Information (NCBI). Reference genomic sequence of HBB gene and neighboring sequences were obtained from NCBI gene database. Using SnapGene desktop tool, exact site of rs334 was located and 100 bp length target sequence centering the possible mutation site was picked. Guide RNA sequences were determined using online gRNA design tool (available at http://crispr.mit.edu/) by Zhang Lab, MIT. 13 candidate gRNA sequences were suggested by the tool, listing the on-target scores and possible off-target numbers for each gRNA. The gRNA sequence which has the highest score and closest distance to the rs334 was picked. Moreover, 7 bp longer version of this gRNA sequence was designed. The gRNA sequence obtained as 20 bp length was elongated from its 5’ direction, resulting in 27 bp long gRNA (**Table 1**).

**Table 1.**
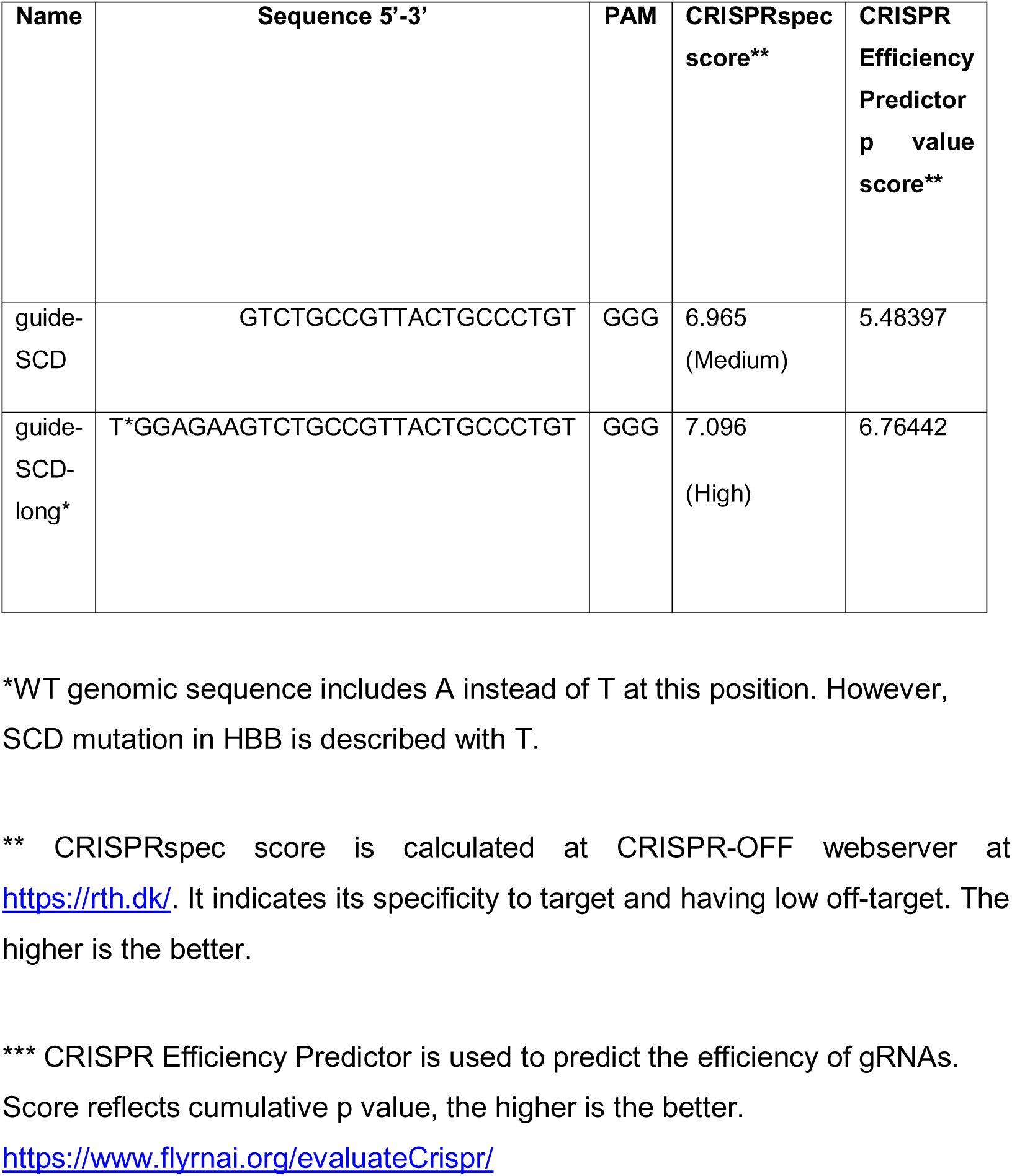
gRNA sequences

### 2.3. Annealing of gRNA sequence

gRNA oligonucleotides were annealed with complementary sequences of the gRNA, which were determined using SnapGene. Adapter sequences (Oligo-1: 5’-CACCGG and Oligo-2: 5’-AAC and C’3) were added to facilitate cloning of designed oligos according to manufacturer’s procedure (REF). Annealing reaction mixture of ssDNA oligos were prepared in PCR tube as Forward Oligo (100 µM) 1 µl, Reverse Oligo (100 µM) 1 µl, 5X Rapid Ligation Buffer 2 µl, T4 Polynucleotide Kinase 1 µl, and Nucleotide Free Water 5 µl as we have done previously (Kocabas et al., 2015a). Prepared reactions were placed into the thermocycler and temperatures were set as 37°C for 30 minutes, 95°C for 5 minutes, Ramp down from 95 to 25°C with 0.1°C /sec cooling. Afterwards, annealed oligos were diluted 1:200 using nuclease free water and stored in - 20°C until use.

### 2.4. Digestion and isolation of the linearized pSpCas9(BB)-2A-GFP

Digestion of pSpCas9(BB)-2A-GFP plasmid by BpiI enzyme was conducted by setting up the reaction mix as 2 µl of 10X FastDigest green buffer, plasmid DNA up to 5 µg, 1 µl of BpiI FastDigest Enzyme, and completed to 20 µl with nuclease free water as we have done previously (Kocabas and Aslan, 2015; Kocabas et al., 2012a; Kocabas et al., 2015a; Kocabas et al., 2015b; Mahmoud et al., 2013; Rimmele et al., 2015; Simsek et al., 2010). Reaction mix was placed in thermocycler for 30 minutes at 37°C for incubation. Following incubation, FastAP Thermosensitive Alkaline Phosphatase was added into the digestion mix as 4 µl of Alkaline Phosphatase, 3 µl of 10X FastAP Buffer, and 3 µl of nuclease free water. Tube was placed back into the thermocycler and set to 37°C for 20 minutes followed by 80°C for 15 minutes. This is followed by DNA gel purification. 1%(w/v) agarose gel was prepared in TBE buffer. Digested and dephosphorylated plasmid mix was loaded into the well. Electroporation was conducted for 40 minutes at 150 V. After 40 minutes, agarose gel was visualized under UV transilluminator. Band around 9.2 kb was dissected using a scalpel and placed into eppendorf tube, and the weight was measured. NucleoSpin® Gel and PCR Clean-up Kit was used for plasmid purification from the gel. To solubilize the gel slice, 200 µl of NTI buffer was added for each 100 mg of agarose gel. Sample was incubated for 5-10 minutes at 50°C until the gel was dissolved, and vortexed briefly. Column was placed into a collection tube and up to 700 µl of sample was loaded. Sample was centrifugated at 11,000 x g for 30 seconds. Flow-through was discarded. To wash the silica membrane, 700 µl of NT3 Buffer was added into the column and centrifugated for 30 seconds at 11,000 x g. Flow-through was discarded. Column was centrifugated at 11,000 x g for 1 minute to dry the membrane. Column was placed into a clean eppendorf tube and 30 µl of prewarmed NE buffer was added directly over the membrane. Sample was incubated at room temperature for 1 minute, then centrifugated for additional 1 minute at 11,000 x g. Concentration was measured using NanoDrop.

#### 2.4.1. Ligation of the pSpCas9(BB)-2A-GFP Plasmid Backbone and gRNA Insert

Ligation reaction was set as 50 ng of purified pSpCas9(BB)-2A-GFP plasmid backbone, 1 µl of annealed gRNA Oligos (1:200), 1 µl of T4 DNA Ligase, 2 µl of 10X T4 DNA Ligase Buffer, and completed up to 2u µl with nuclease free water as we have done previously (Kocabas and Aslan, 2015; Kocabas et al., 2012a; Kocabas et al., 2015a; Kocabas et al., 2015b; Mahmoud et al., 2013; Rimmele et al., 2015; Simsek et al., 2010). Reaction mix was incubated in thermocycler at 22°C for 90 minutes. Tubes were left overnight at RT. Mix was used for bacterial transformation by heat shock procedure.

### 2.5. Bacterial Transformation by Heat Shock

Chemically competent DH5α cells (50 µl) were thaw on ice. Ligated plasmids were placed on ice for 10 minutes. 7 µl of ligation mix was added into bacteria by pipetting gently. Bacteria mixture was incubated on ice for 25 minutes. Tubes were placed on heat block adjusted to 42°C for 45 seconds and immediately placed on ice again for 3 minutes to provide heat shock. 250 µl of SOC medium was added to each tube. Cells were incubated in shaking incubator at 37°C, 150 rpm for 120 minutes. Inoculum was spread over Ampicillin (+) agar plates. Plates were incubated overnight in 37°C incubator. Colonies were observed the next day. This is followed by colony selection and miniprep isolation from individual colonies as we have done previously (Kocabas and Aslan, 2015; Kocabas et al., 2012a; Kocabas et al., 2015a; Kocabas et al., 2015b; Mahmoud et al., 2013; Rimmele et al., 2015; Simsek et al., 2010).

### 2.6. Verification of gRNA insert by Polymerase Chain Reaction (PCR)

The primers listed in the **Table 2** were used for PCR amplification to test the presence of gRNA inserts on plasmids. The primer for Cloning Forward was common for both gRNA inserted plasmids, annealing U6 promoter upstream of the gRNA cloning site. For each gRNA, original reverse oligos used annealing of gRNA were used as reverse primers for PCR reaction. Stock primers (100 µM) were diluted to 1:20 using nuclease free water. PCR reaction was set according to the instructions. 10 µl of 2X Master Mix, 1 µl of Plasmid DNA, 1 µl of Forward Primer and Reverse Primer, 7 µl of Nuclease Free Water. Thermocycler was adjusted as Initial denaturation at 95°C for 60 sec, 25 cycles of (Denaturation at 95°C for 15 sec 25 cycles, and Annealing at 55°C for 15 sec, Extension at 68°C for 10 sec), and final extension at 68°C for 2 minutes. 2% agarose gel was prepared using TBE buffer. Into each PCR product, 4 µl of 6X loading dye was added. 4 µl of 50 bp DNA Step Ladder was loaded into the first well. PCR products were loaded as 10 µl in each well. Gel was run at 150 V for 35 minutes. Gel image was obtained from Biorad Chemidoc XRS device.

**Table 2.**
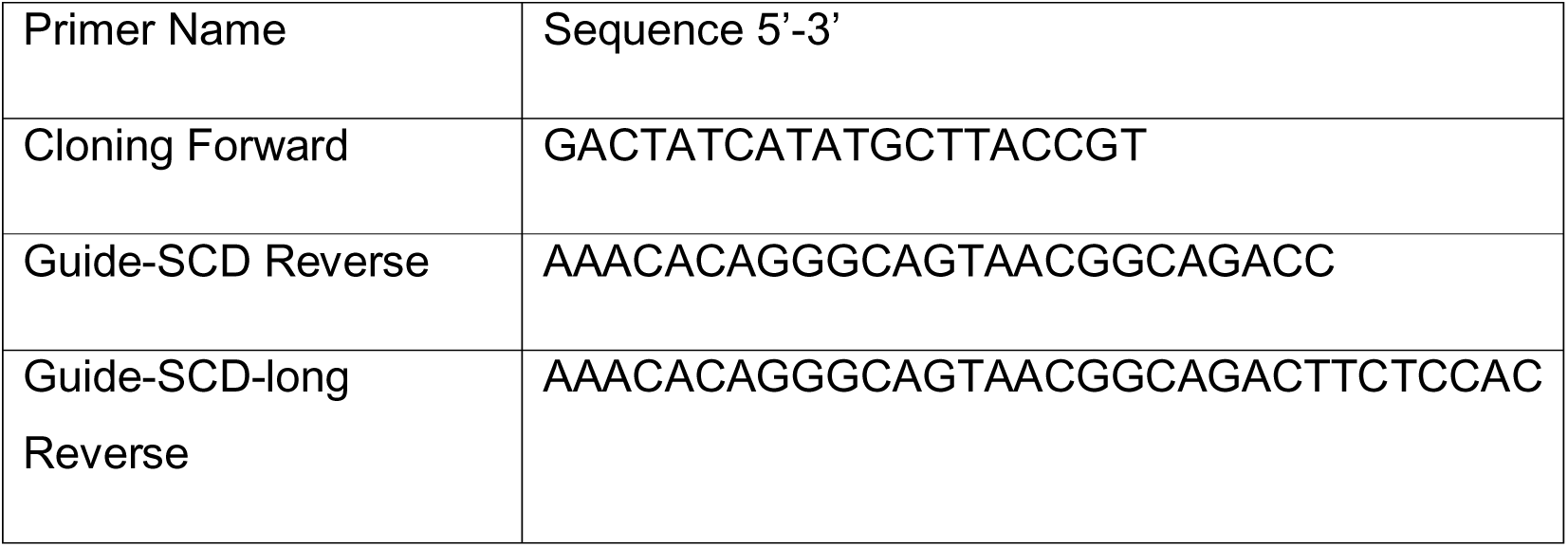
Primer sequences for PCR reaction

### 2.7. Verification of the gRNA insert by Sanger Sequencing

Plasmids isolated by midiprep protocol were sent to Medsantek, Turkey for sequencing. Directional Sanger sequencing was performed using 10 µM primer Cloning Forward given in **Table 2** was used. The plasmid DNA concentrations were adjusted to 100 ng/µl.

### 2.8. Mammalian cell transfections

Polyethylenimine (PEI) was used for mammalian cell transfections as we have done previously (Boztas et al., 2013; Kocabas and Aslan, 2015; Kocabas et al., 2015a; Rimmele et al., 2015). Briefly, HEK293T cells were seeded in 6 well plate with 3×10^5^ density and cultured in 2 ml DMEM high glucose supplemented by 10% FBS, no antibiotics. Cells were incubated overnight at 37°C, 5% CO_2_ and 95% RH. In a 1.5 ml tube, 1 µg pSpCas9(BB)-2A-GFP-gRNA was diluted in 200 µl serum-free DMEM. Control mix was prepared with pSpCas9(BB)-2A-GFP without gRNA. 2 µg PEI (1 mg/ml stock) was added and mixed well (DNA: PEI ratio is 1:2). The mixture was incubated at room temperature for 15 minutes. The mixture was added to the culture media dropwise and mixed without disturbing the cells. Cells were placed in the incubator and incubated for 16-24 hours. GFP expression was checked under fluorescent microscope. GFP expression and confluency was analyzed. Cells were harvested by trypsinization for downstream applications. Transfection efficiency was analyzed by fluorescence microscopy and flow cytometer based on the analysis of GFP. To this end, 24 hours after transfection, cells were examined under inverted fluorescence microscope (Zeiss Axio Vert.A1) as we have done previously (Mahmoud et al., 2013). Filter was set to FITC mode and imaging was performed with 5X and 10X magnifications. Images were recorded in TIFF format. Flow cytometer were used to determine GFP expressing cell population using Beckman Coulter’s CytoFLEX S flow cytometry device as we have done previously (Aksoz et al., 2018; Kocabas et al., 2015b; Kocabas et al., 2012b; Simsek et al., 2010; Zheng et al., 2014). HEK293T cells were harvested by trypsin and washed twice with PBS. Cell suspensions were transferred into 96 well plates and the plates were placed to the plate reader part of the device. Acquisition was set to 1 x10^5^ events read in live population. Live population was determined according to distinct aggregation on forward to side scatter plot. Gate was taken covering live population and GFP expression was analyzed at FITC-A / FSC-A dot plot. Non-transfected cells were used as GFP negative control. Results were recorded as histogram and dot plot.

### 2.9. Detection of indels by standard and long gRNAs

Genomic DNA was isolated from cell transfected with pSpCas9(BB)-2A-GFP, or pSpCas9(BB)-2A-GFP vectors containing standard gRNA or long-gRNA. Cells were resuspended in 200 µl PBS. 20 µl of Protainase K were added into the sample followed by addition of 20 µl of RNase A and mixed well via vortex and incubated at RT for 2 minutes. 200 µl of Lysis/Binding Buffer was added and mixed by vortexing. Sample was kept at 55°C for 10 minutes. 200 µl of absolute ethanol was added to the lysate and mixed well by vortexing for 5 seconds. Lysate was transferred into spin column. The column was centrifuged at 10.000 x g for 1 minute. Collection tube was discarded, and column was placed into a new collection tube. 500 µl of Wash Buffer 1 was added to the column and centrifuged at 10.000 x g for 1 minute. Collection tube was replaced and 500 µl of Wash Buffer 2 was added to the column. Column was centrifuged at maximum speed for 3 minutes. Collection tube was discarded, and column was placed in a 1.5 Eppendorf tube. 50 µl of Genomic Elution Buffer was added to the column and incubated at RT for 1 minute. Tube with column was centrifuged at max speed for 1 minute. Elution step was repeated for maximum recovery of DNA. Column was discarded and DNA within the tube was measured with NanoDrop. This is followed by PCR amplification of target locus.

The primers listed in the **Table 3** were used for PCR amplification of endonuclease targeted locus of HBB gene. Primers were designed to produce 1 kb products centering the expected target site. EnGen™ Mutation Detection Kit was used for PCR amplification. PCR reaction was done with Q5 Hot Start High-Fidelity 2X Master Mix according to manufactures recommendations.

**Table 3.**
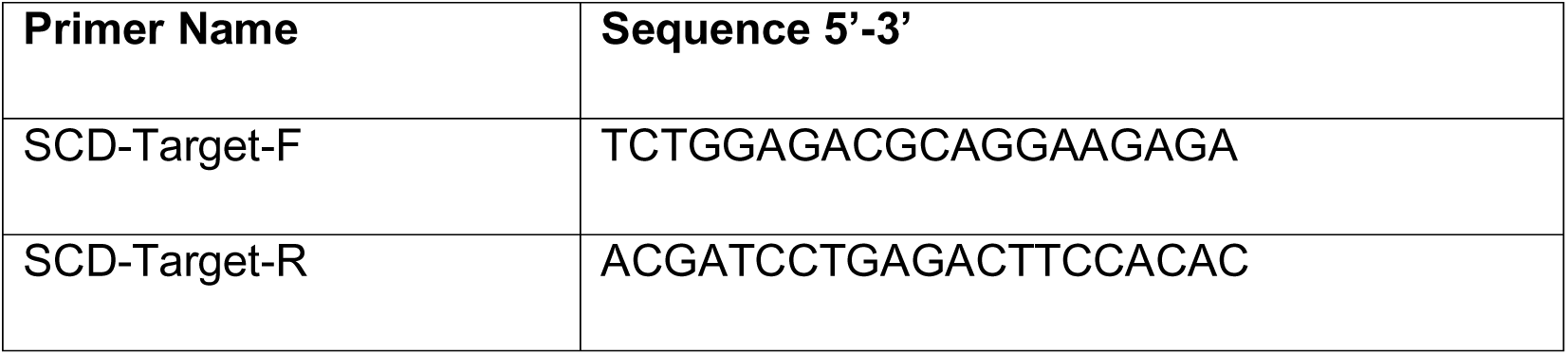
PCR Primer sequences for HBB loci

Annealing temperatures for the primers were found at https://tmcalculator.neb.com/ website and reaction were set on thermocycler accordingly. Thermocycler was adjusted as initial denaturation at 98°C for 30 sec, 25 cycles of (denaturation at 98°C for 7 sec 35 cycles, and annealing at 55°C for 12 sec, Extension at 72°C for 20 sec), and final extension at 72°C for 2 minutes.

Thermocycler was also set for the control reaction according to manufactures recommendations.: After the reactions, 5 µl of PCR product was spared for further applications and remaining was mixed with 1 µl Purple Gel Loading dye (6X). 1% agarose gel was prepared with TBE buffer and samples were loaded. Gel was run at 150 V for 45 minutes. Gel image was obtained from Biorad Chemidoc XRS device.

#### 2.9.1. T7 Endonuclease Assay

Thermocycler was used to denature and anneal 5 µl of PCR product as recommended by manufacturer (EnGen™ Mutation Detection Kit, NEB, E3321S). Thermocycler was adjusted for heteroduplex formation as following: Denaturation at 95°C for 5 minutes, annealing at 95-85°C at 1.5°C/min ramp rate for 5 sec, annealing at 85-25°C at 0.1°C/min ramp rate for 5 sec and Hold at 4°C until use. To digest the heteroduplex, 1 µl T7 Endonuclease I was added into the mixture and incubated for 20 minutes at 37°C in the thermocycler. Afterwards, 1 µl of Proteinase K was added to inhibit T7E activity and incubated 37°C for 5 minutes in the thermocycler. Sample was mixed with 4 µl of Purple Gel Loading dye (6X) and loaded on 1% agarose gel prepared with TBE Buffer and run at 150 V for 45 minutes. Gel image was obtained from Biorad Chemidoc XRS device. ImageJ was used to quantify each product with band density function as we have done previously (Mahmoud et al., 2013) to determine % of indels. To this end, digested band densities and non-digested band intensities quantified. Digested band density divided by total band density to determine the percentage of indel formation.

### 2.10. PCR Amplification and TA cloning of wild type genomic DNA HDR

The primers listed in the **Table 4** were used for PCR amplification of wild type genomic DNA sequence of HBB gene. Primers were designed to produce 1 Kb and 2 Kb long products centering the expected target site. PCR reaction was prepared using *Taq* 2X Master Mix to create 2 different PCR product lengths as 1 Kb, and 2 Kb. Reaction was prepared as recommended by manufacturer. This is followed by TA cloning of PCR products into pCRv2.1 Vector with T4 DNA Ligase according to manufactures procedures (TA Cloning™ Kit, Invitrogen, K202020). Ligation reaction was prepared and incubated at RT for 15 minutes as we have done previously (Aksoz et al., 2018; Kocabas et al., 2015b; Kocabas et al., 2012b; Simsek et al., 2010; Zheng et al., 2014). Positive clones were selected based on Blue-White Screening by X-gal staining. Transformed bacteria were spread over agar plate containing 100 µg/ml Kanamycin, 40 µl of 40 mg/ml X-Gal and 40 µl of 100 mM IPTG. Plates were incubated at 37°C overnight and observed the next day. Following miniprep DNA isolation of white colonies, plasmids were digested with EcoRI. Reaction was incubated at 37°C for 1 hour. Sample was mixed with 4 µl of Purple Gel Loading dye (6X) and loaded on 1% agarose gel prepared with TBE Buffer and run at 150 V for 45 minutes. Gel image was obtained from Biorad Chemidoc XRS device.

**Table 4.**
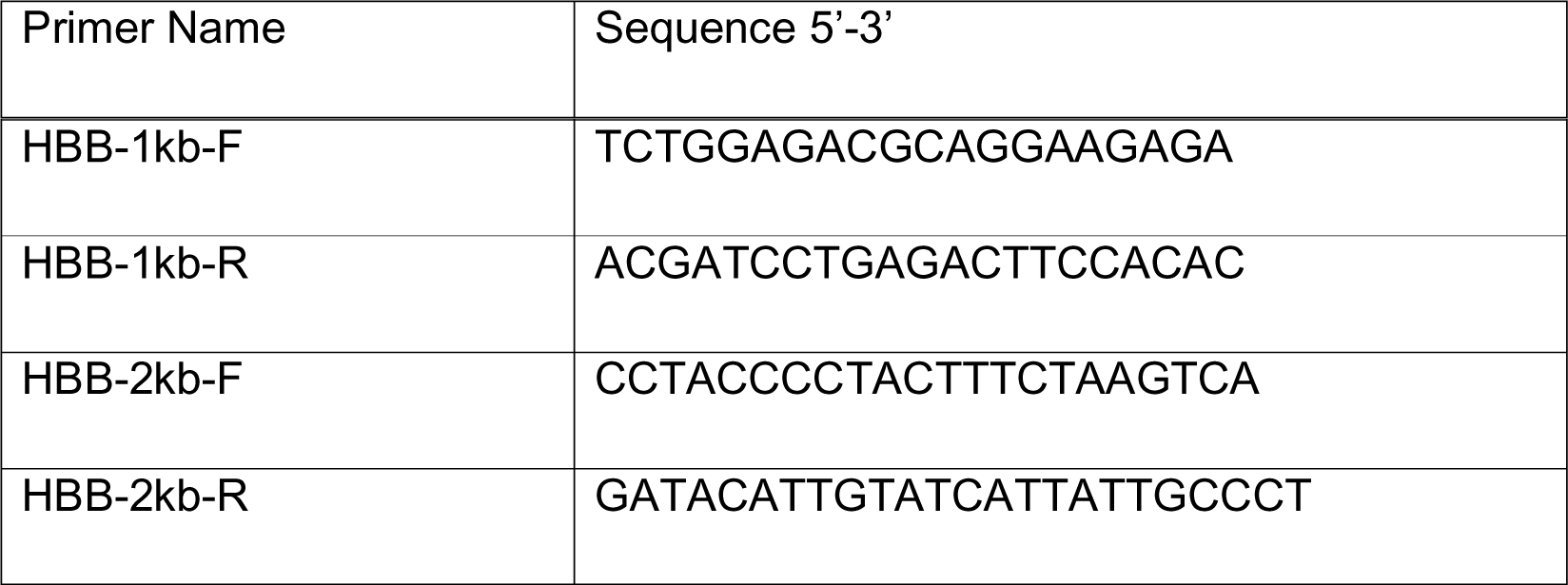
Primer sequences for PCR amplification of wild type genomic DNA

### 2.11. Site Directed Mutagenesis in HDR template

Primers for site directed mutagenesis were designed using online In-Fusion Cloning Primer Design Tool (available https://www.takarabio.com/learning-centers/cloning/in-fusion-cloning-tools). The primers listed in the **Table 5** were used for plasmid linearization and induction of sequence changes. Sequences included HindIII for downstream verification and removal of PAM sequence for repetitive Cas9 binding and induction of DSBs. PCR mix was prepared using the plasmids obtained from TA cloning and CloneAmp HiFi PCR Master Mix according to manufactures recommendations (In-Fusion® HD Cloning Plus, Takara, 638910). This is followed by in-fusion reaction for site directed mutagenesis as we have done previously (Kocabas and Aslan, 2015; Kocabas et al., 2015a). Reaction was incubated at 50°C for 15 minutes. Plasmids were digested using HindIII restriction enzyme for verification of site directed mutagenesis and run 1% agarose gel. ImageJ was used to determine (Mahmoud et al., 2013) percent of HDR event by analyzing corresponding band densities, and by normalizing the transfection efficiency as determined by flow cytometric analysis of GFP+ cells.

**Table 5.**
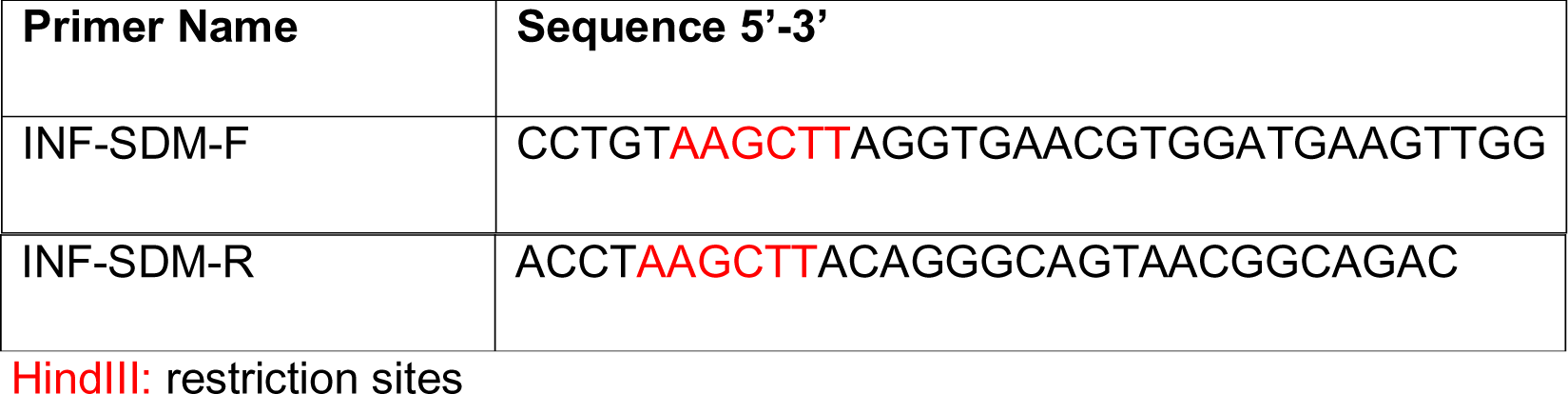
Primer sequences for inverse PCR reaction to be used for in fusion cloning

### 2.12. Transfection of HEK293T Cell Line With spCas9(BB)-2A-GFP gRNA vectors and HDR Templates

Transfection HEK293T cells with HDR templates and spCas9(BB)-2A-GFP gRNA vectors was performed in various combinations shown in **Table 6** with PEI method. 12 hours after transfection, culture media was changed with fresh complete DMEM supplemented by SCR7 with final concentration of 1µM. PCR Amplification of Donor Template post gene targeting was performed. The primers HBB-1kb-F (5’-TCTGGAGACGCAGGAAGAGA-3’) and HBB-1kb-R (5’-ACGATCCTGAGACTTCCACAC-3’) were used for PCR amplification endonuclease targeted genomic DNA sequence of HBB gene. Primers were designed to produce 1 kb long products centering the expected target site. PCR product obtained was digested using HindIII restriction enzyme. Briefly, reaction mixture was placed in thermocycler for 15 minutes at 37°C. Afterwards, products were loaded on 1% agarose gel and run at 150 V for 1 hour. Results are obtained from Biorad Chemidoc XRS device.

**Table 6.** Mixtures for transformation via PEI

| TUBE # | spCas9(BB)-2A-GFP vectors | 1 kb donor | PEI | TUBE # | spCas9(BB)-2A-GFP vectors | 2 kb donor | PEI |
| --- | --- | --- | --- | --- | --- | --- | --- |
| 1 | 0 µg | 1 µg | 2 µg | 7 | 0 µg | 1 µg | 2 µg |
| 2 | 1 µg | 0 µg | 2 µg | 8 | 1 µg | 0 µg | 2 µg |
| 3 | 0,5 µg | 0,5 µg | 2 µg | 9 | 0,5 µg | 0,5 µg | 2 µg |
| 4 | 0,5 µg | 1 µg | 3 µg | 10 | 0,5 µg | 1 µg | 3 µg |
| 5 | 1 µg | 1 µg | 4 µg | 11 | 1 µg | 1 µg | 4 µg |
| 6 | 1 µg | 2 µg | 6 µg | 12 | 1 µg | 2 µg | 6 µg |

## 3. Results

### 3.1. Verification of gRNA inserts into pSpCas9(BB)-2A-GFP vector

To confirm the successful cloning of gRNA sequences into pSpCas9(BB)-2A-GFP plasmids, PCR amplification for the cloning site was performed. In PCR reaction, the primer named Cloning-F was used as the standard forward primer annealing the U6 promoter located upstream of the gRNA cloning site (**Figure 1A**). The reverse strands of oligo DNAs previously designed and used for gRNA insert production were used as the reverse primer for each gRNA inserted plasmid. It was expected to obtain a 100 bp PCR product if the gRNA sequence is successfully cloned in to plasmid. In the absence of gRNA insertion, reverse primer would not anneal to template plasmid DNA therefore there would not be any visualized PCR product in agarose gel. Cloning results were also verified with Sanger sequencing method. PCR reactions run on 1% (w/v) agarose gel to assess successful cloning of gRNA oligos into into pSpCas9(BB)-2A-GFP vector (**Figure 1B**). PCR result indicates successful cloning process of two gRNA sequences, standard HBB gRNA and SCD specific HBB long gRNA, into pSpCas9(BB)-2A-GFP. The plasmid without any gRNA insert did not produce any 100 bp PCR products while gRNA cloned plasmids did, indicating the presence of gRNA sequences downstream of the U6 promoter. In addition, Sanger sequencing of gRNA plasmids have shown that both gRNA sequences were successfully cloned into plasmids and ready for downstream applications (**Figure 1C**). Locations of each insert are at the correct sites to facilitate gRNA and gRNA scaffold expressions in transfected cells.

**Figure 1.**
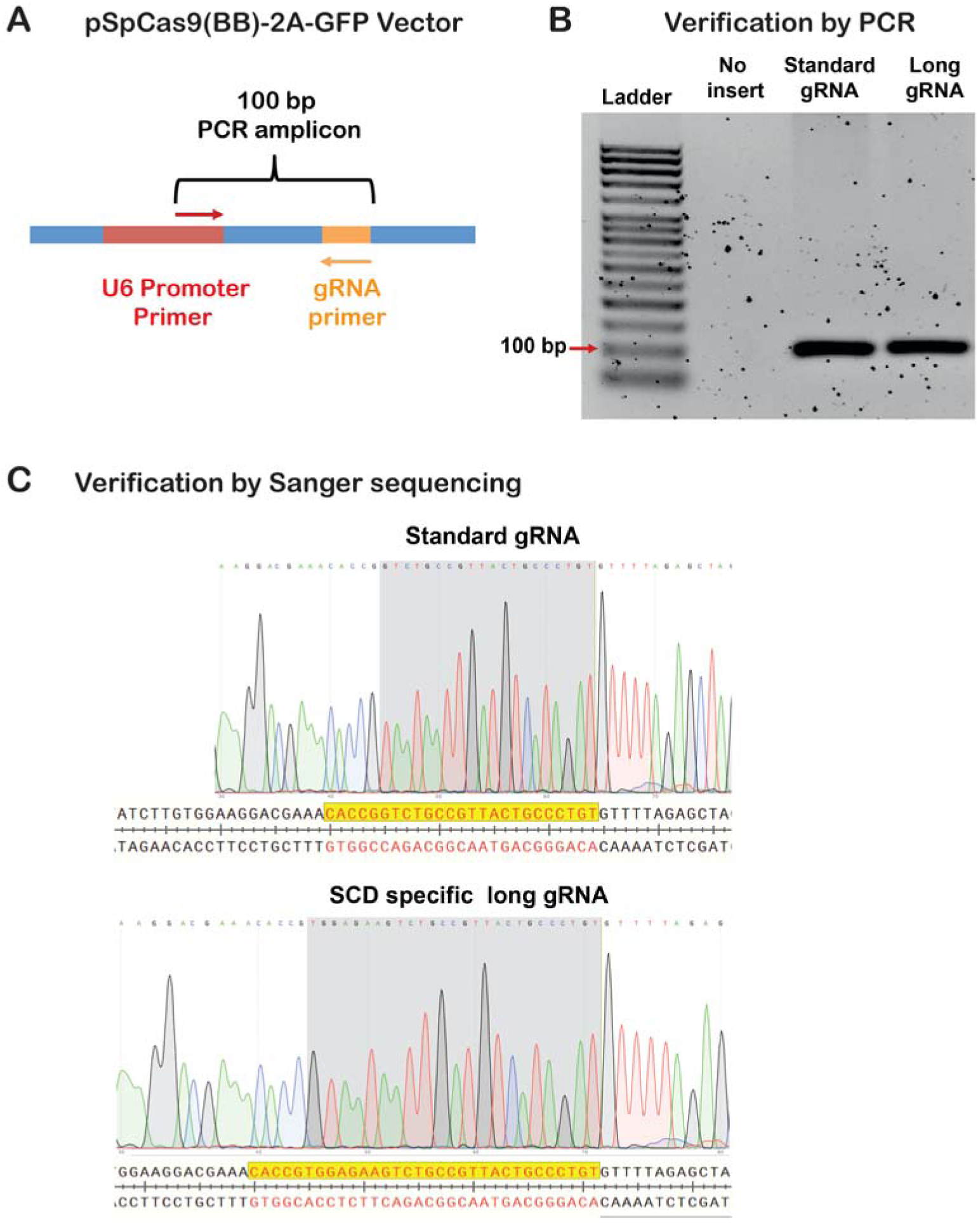
Verification of standard gRNA and SCD specific long-gRNAs. **A)** Schematic of PCR primers designed to verify cloning of gRNAs into pSpCas9(BB)-2A-GFP vector. U6 promoter primer and gRNA oligos were used. **B)** Verification of gRNAs cloning with PCR. Agarose gel image of PCR products for no insert (empty vector), standard gRNA cloned vector, and long-gRNA cloned vector. **C)** Verification of gRNA sequences in pSpCas9(BB)-2A-GFP vector by Sanger sequencing.

### 3.2. Detection of Indel formation at HBB locus

Equal number of cultured HEK293T cells were transfected with the same amount of pSpCas9(BB)-2A-GFP plasmids using PEI transfection reagents to figure out the most efficient way to deliver Cas9 endonuclease, gRNA and/or HDR templates. 24 hours post-transfection, cells were examined under fluorescent microscope (**Figure S1**). Each well containing 300,000 cells were transfected with 1 µg of plasmid. In PEI transfections, to obtain 1:2 and 1:3 DNA:PEI ratios, 2 µg and 3 µg of PEI were used respectively (Figure S1A). PEI has shown optimal transfection efficiency and GFP expression at these ratios. The increased amount of PEI resulted in an increase in cell death. Thus, we have used 1:2 DNA to PEI ratio in transfection studies in HEK293T cells. PEI transfection of pSpCas9(BB)-2A-GFP vectors (1:2) was repeated for HEK293T cells to assess transfection efficacy. Cells were harvested 24 hours after transfection by trypsinization and GFP expression was analyzed using flow cytometry (Figure S1B). pSpCas9(BB)-2A-GFP vectors containing standard HBB gRNA and HBB long-gRNA showed 40-50% transfection efficiencies.

T7 Endonuclease (T7E) based assay was used to evaluate the activity of both gRNA on target locus. Briefly, genomic DNAs were isolated from HEK293T cells transfected with HBB gRNA and HBB long-gRNA Cas9-GFP. Primers for the gDNA PCR were designed to produce 1 kb product, centering the DNA sequence targeted by gRNAs. It was expected to generate 500 bp band on agarose gel after T7E treatment to PCR products if the gRNA successfully induce DSBs resulting in indel mutations. PCR products derived from gene targeted cells were proceeded to mismatch cleavage assay, followed by agarose gel electrophoresis (**Figure 2A**). The data indicates that both HBB gRNA and HBB long-gRNA pSpCas9(BB)-2A-GFP vectors successfully functioned and showed similar cleavage activity (up to 40 % indels in whole cells) on target DNA sequence (**Figure 2B**). Cells transfected with Cas9 only did not show any DSB on target locus since there was not a gRNA for gene targeting. In addition, when we normalized the % indels per GFP+ cells (see Figure S1C), we have found that studied gRNAs are able to cause indel formation up to 88% in transfected cells (**Figure 2C**). Intriguingly, longer gRNA has showed slightly better performance of targeting and cleavage on target locus, in comparison to standard length gRNA. In theory, longer gRNA would have less off-target effects on genome. Since long gRNA has the same or better targeting efficiency and functionality on target site, it could be preferred to use for future application rather than using standard length gRNA. Thus, it would be possible to reduce off-target mutations without losing on-target activity. Indeed, off-target predictions demonsrated a significant reduction in the off-targets by long gRNA targeting SCD mutation. HBB standard gRNA and long gRNA were subjected to off-target prediction with Cas-OFFinder tool. Numbers of off-target were compared for every given parameter. DNA bulge, RNA bulge 0,1 and 2 were characterized along with upto 9 mismatches to determine potential targets. We have determined up to 99% reduction by HBB long gRNA compared to standard gRNA off-targets predicted (**Supplementary File 1**).

**Figure 2.**
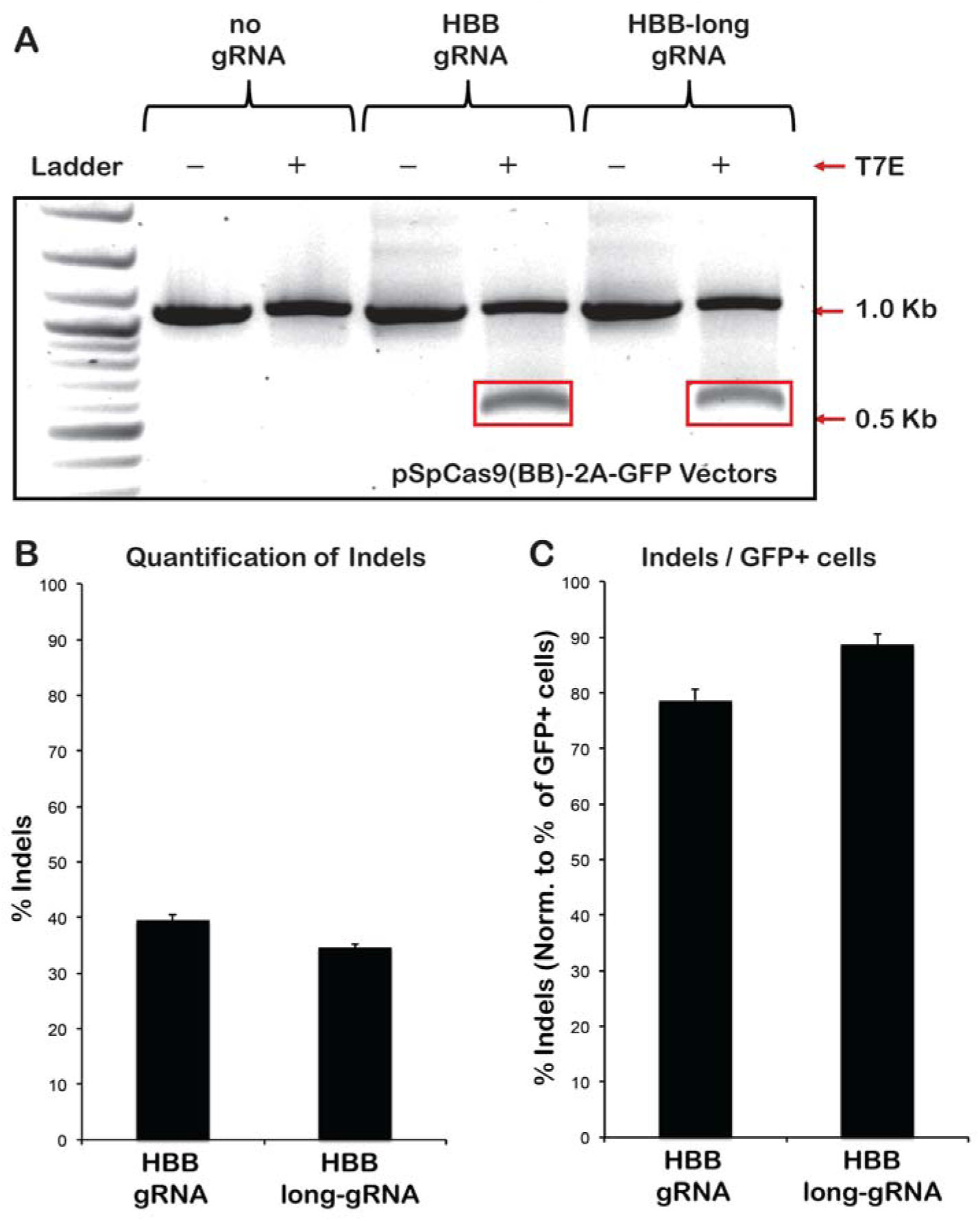
Mismatch cleavage assay. **A)** T7 endonuclease assay was used to assess heteroduplex formation by indels formed by gRNA vectors transfected into HEK293T cells. Cells were transfected with pSpCas9(BB)-2A-GFP only, standard size HBB-gRNA pSpCas9(BB)-2A-GFP vector, and HBB-long gRNA pSpCas9(BB)-2A-GFP vectors. PCR amplicons were in 1 Kb size. Note that gRNAs targeted the middle of the PCR amplicons. **B)** Quantification of percentage of Indels formed by gRNAs. **C)** Quantification of percentage of Indels formed per GFP+ cells. Note that band intensities in the red boxes were quantified and normalized to total in the corresponding lane. Note that we obtained upto 88% indel formation with HBB long-gRNA.

### 3.3. Generation of HDR templates

Wild type genomic DNA from HEK293T cells was isolated and target locus encompassing HBB gene was amplified by PCR to perform TA cloning (**Figure 3A**). Taq polymerase was used to obtain PCR products with TA overhangs. Primer pairs were designed to produce 1kb, and 2kb DNA fragments centering the gRNA binding site. PCR reactions were prepared, and product sizes were examined by agarose gel electrophoresis (**Figure 3B**). The result indicates that 1 Kb and 2 Kb fragments were successfully produced in correct size and without any unspecific products. PCR products with 1 Kb and 2 Kb size were gel purified and were directly used for TA cloning protocol. DNA fragments were ligated to linearized pCR 2.1 plasmid and bacterial transformation was performed. The cloning site of this plasmid is in LacZ alpha sequence, allowed us to perform the blue-white screening of transformed colonies. White colonies were selected and plasmids were isolated. To ensure the insert size in the selected plasmids, EcoRI digestion was performed. Digestion with EcoRI confirmed the correct insertions into pCR 2.1 plasmids (**Figure 3C**, colony 2 for 1 Kb HDR amplicon and colony 1 for 2 Kb HDR amplicon). Plasmids with about 4.9 Kb and 5.9 Kb total size were selected for the site directed mutagenesis step to replace PAM sequence with HindIII restriction site.

**Figure 3.**
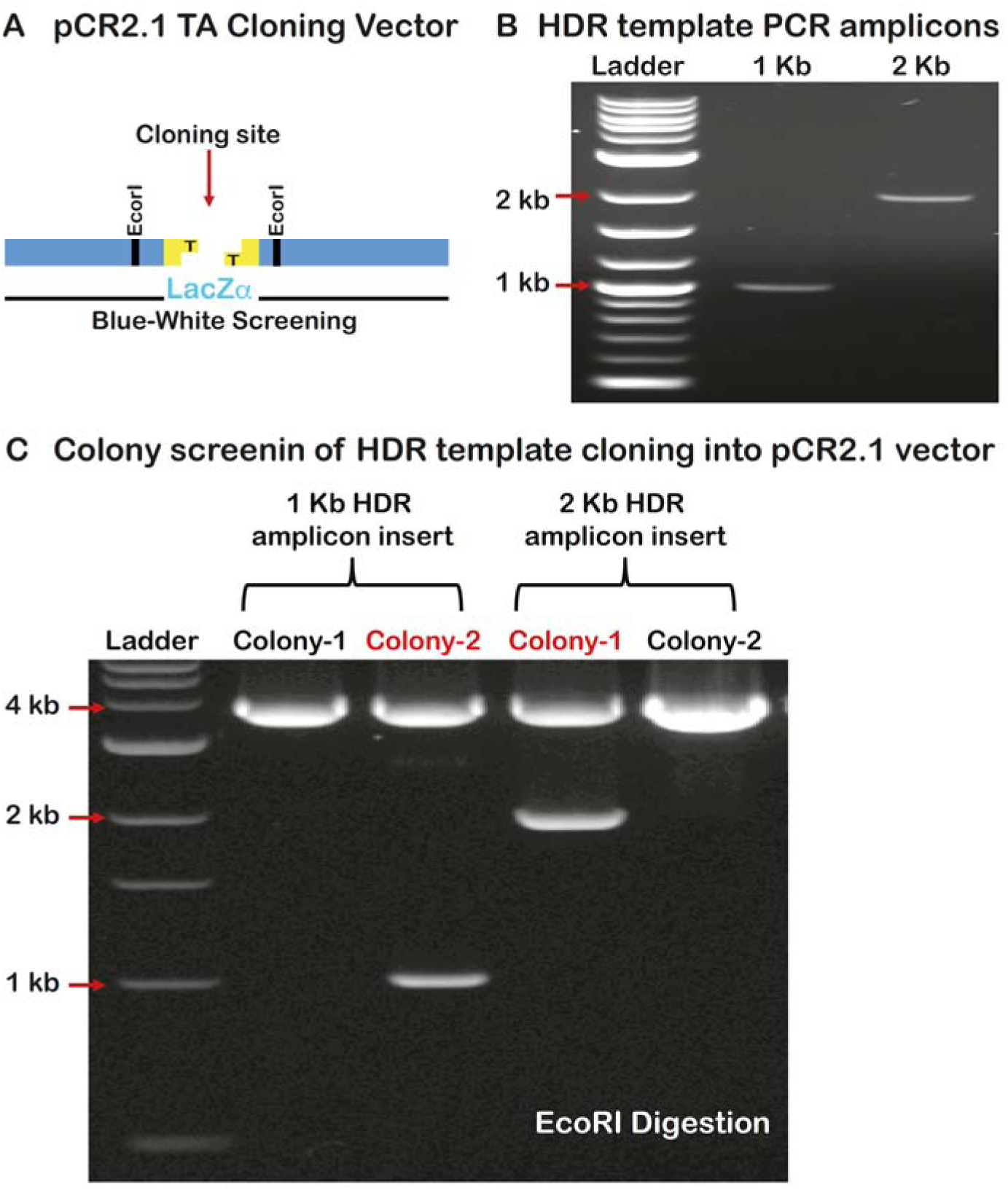
HBB HDR repair template generation. **A)** Schematic of TA cloning into pCR2.1 vector. Agarose gel images demonstrate **B)** PCR amplification of HBB 1Kb and 2 Kb HDR repair templates and **C)** Verification of cloning into pCR2.1 vectors with EcoRI digestions. At least 2 colonies were screened for each.

We aimed to achieve targeted gene editing of SCD mutation to WT sequence via HDR mechanism. In addition, PAM sequence on the wild type sequence aimed to be replaced with HindIII restriction enzyme recognition sequence (**Figure 3A**). To this end, we used in-fusion cloning system to mutagenize and generate HBB-HDR-1Kb-NoPAM (**Figure 3B**) and HBB-HDR-2Kb-NoPAM (**Figure 3C**) plasmids. To confirm the sequence modification on HDR templates, restriction digestion with HindIII enzyme was performed. Digested products were run 1% (w/v) agarose gel. The agarose gel image indicates that the PAM sequence in 1 Kb or 2 Kb wild type HDR templates was successfully replaced with HindIII recognition site (**Figure 4D** **and** 4E).

**Figure 4.**
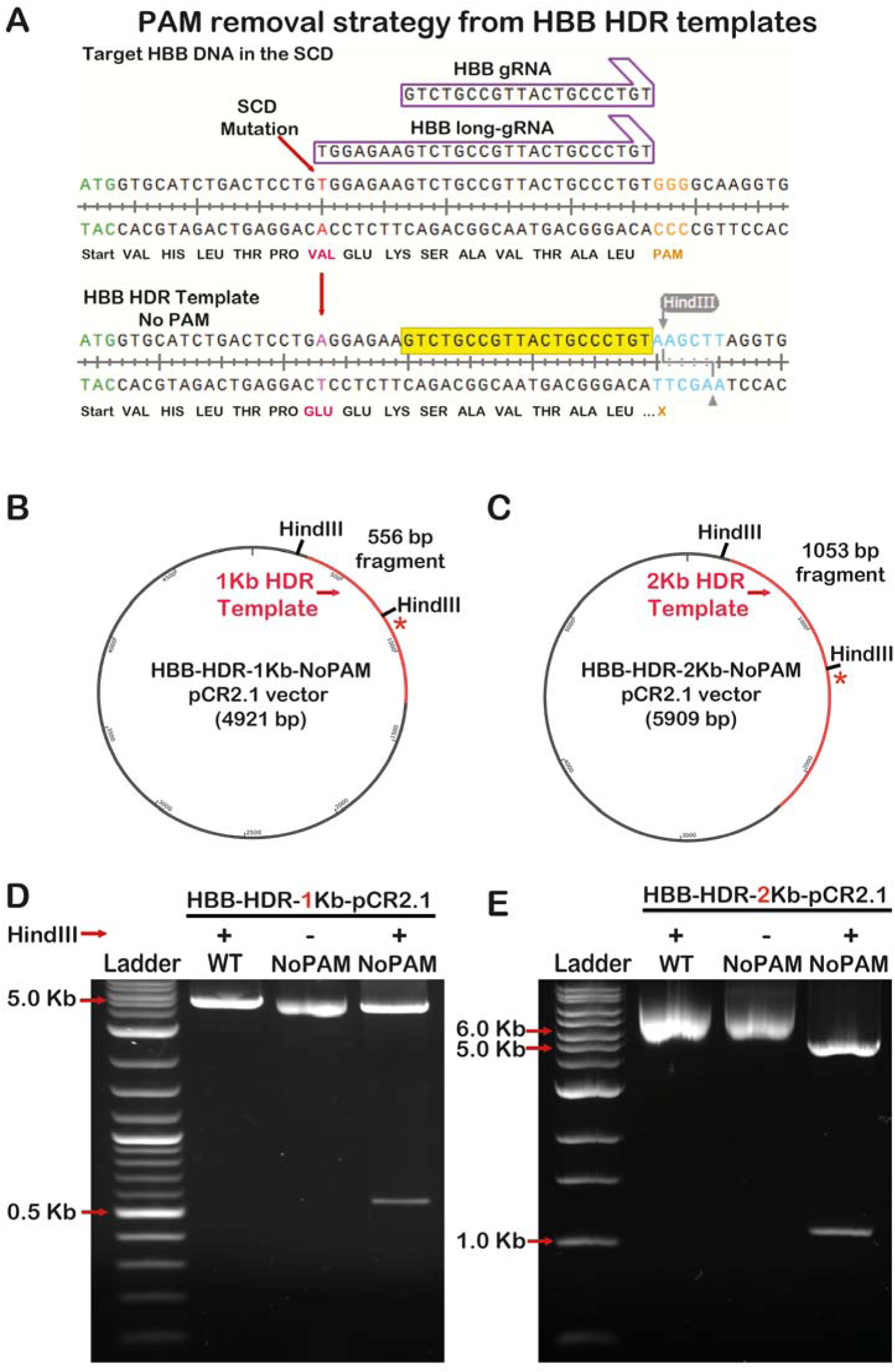
Removal of PAM sequence in the HDR templates by site directed mutagenesis. **A)** Top: Target HBB DNA segment in the genome of SCD patients with tested standard and long gRNAs targeting close proximity of SCD mutation. Bottom: HBB HDR template that replaces PAM sequence with HindIII cut site. Note that this is a proof-of-concept design to remove PAM sequence to prevent re-cutting by spCas9 post HDR template integration, not intended for use as HDR repair template. Plasmids maps of mutagenized **B)** HBB-HDR-1Kb-NoPAM pCR2.1 vector and **C)** HBB-HDR-2Kb-NoPAM pCR2.1 vector. HindIII recognition sequences were introduced in the center to replace PAM sequences. “**\*”** sign shows the where the PAM sequence replaced with HindIII sequence. Agarose gel images show **D)** HindIII digest of HBB-HDR-1Kb-NoPAM pCR2.1 vector and **E)** HindIII digest of HBB-HDR-2Kb-NoPAM pCR2.1 vector.

### 3.4. Co-Delivery of Cas9-gRNA and HDR vectors

The final and most crucial step of this study was the delivery of Cas9, gRNA and HDR templates to induce site-specific genetic modification. HBB long-gRNA-pSpCas9(BB)-2A-GFP vector was introduced in the form of plasmid. HDR templates were delivered as circular plasmids carrying 1kb and 2 kb total homology lengths and the added HidIII site for the screening of HDR events in target cells. Transfection was done using 1:2 DNA to PEI protocol which was previously suggested as the most efficient method for this cell line (**Figure 5A**). HEK293T cells were co-transfected with different amounts of Cas9/gRNA and donor template plasmids (**Table 6** and **Figure B**). Additionally, cells were treated with 1 µM SCR7, which is a small molecule, reported as an HDR enhancer. After 24 hours of transfection, GFP expression in each sample was assessed by flow cytometry (**Figure S2**). Flow cytometry analysis of GFP expression with HBB long-gRNA-pSpCas9(BB)-2A-GFP and 1 Kb (Figure S2A) or 2 Kb (Figure S2B) HDR template vectors exhibited an efficient and similar transfection yield after 24 hours.

**Figure 5.**
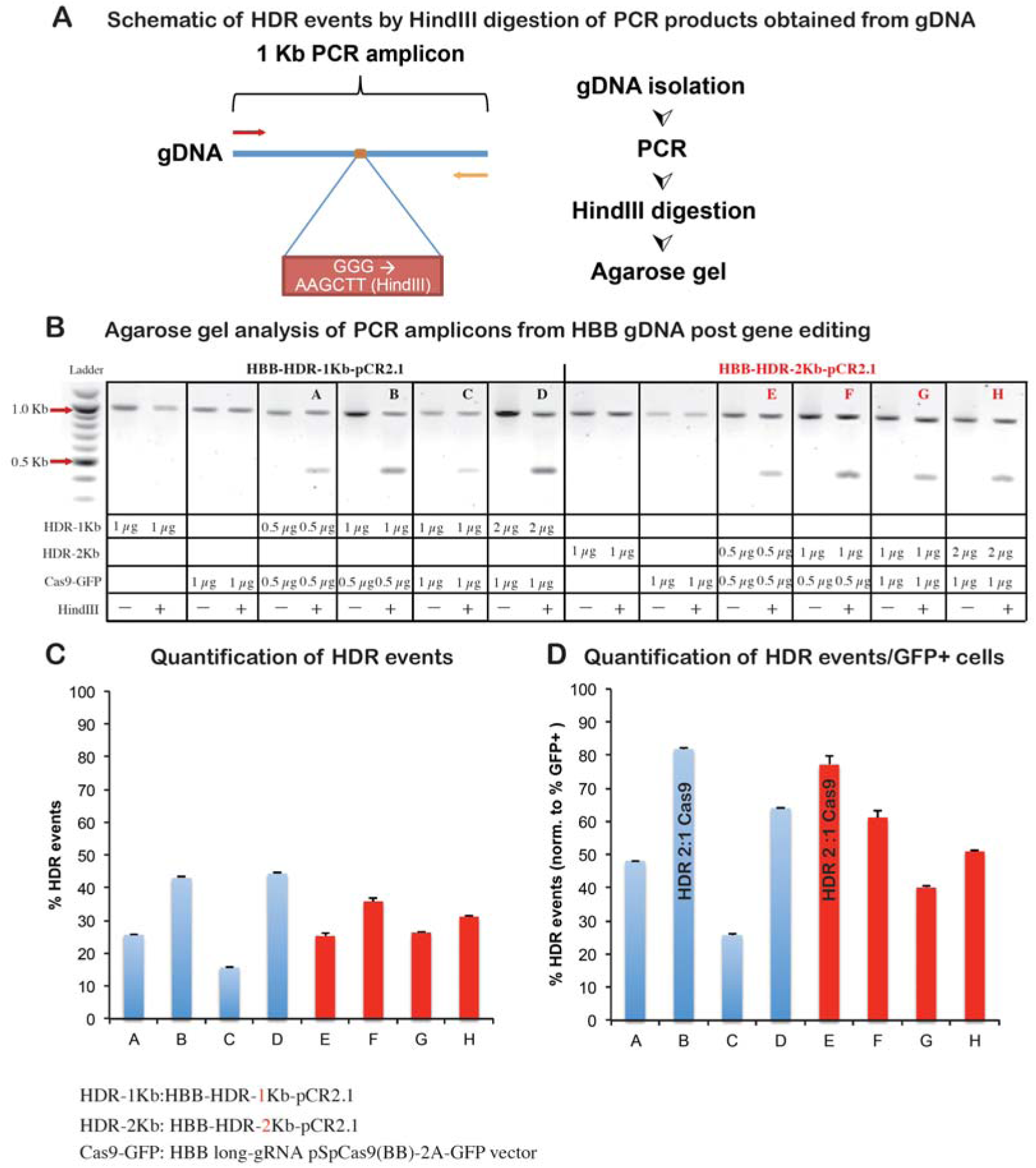
HDR event analysis post transfections. **A)** Schematic of HDR events by HindIII digestion of PCR products obtained from HBB gDNA. **B)** Agarose gel analysis of PCR amplicons from HBB gDNA post gene editing. **C)** Quantification of HDR events. **D)** Quantification of HDR events/GFP+ cells. Note that HDR 2:1 Cas9 ratio yielded highest percentage of HDR event for both 1 Kb and 2 Kb HDR templates. HDR-1Kb: HBB-HDR-1Kb-pCR2.1, HDR-2Kb: HBB-HDR-2Kb-pCR2.1, and Cas9-GFP: HBB long-gRNA pSpCas9(BB)-2A-GFP vector. Note that we obtained up to 80% HDR events with HDR template 2:1 Cas9-GFP transfections.

The efficiency of gene editing on HBB gene, in which the locus where point mutation causing SCD, was examined 72 hours post-transfection and SCR7 treatment. Genomic DNAs were isolated for each sample and the target locus was amplified centering the theoretical location of genomic modification. PCR amplification was followed by HindIII restriction enzyme digestion and the reaction mixture was run on agarose gel (**Figure 5B**). The presence of 0.5 kb band on the gel was the indicator of the replacement of PAM sequence (GGG) with HindIII recognition sequence (AAGCTT) by the HDR mechanisms. Data indicates that the desired genomic modification on HBB gene was achieved with varying efficiencies (**Figure 5C**). When we normalized HDR events per GFP+ transfected cells, we have found up to 80% HDR efficiencies. Intriguingly, the combinations in which the amount of HDR template plasmid was the double of Cas9/gRNA plasmid has shown better efficiency to generate the desired modification on HBB gene. There were not any significant differences between 1kb and 2 kb homology donors in terms of gene editing yield. According to on-target analysis, the required steps were successfully completed to fulfill the mission of sequence alteration introduced by HDR template on HBB gene, with only a few base distances to the point mutation (rs334) causing SCD.

In this study, homology arms were designed symmetrically, 0.5 Kb and 1 Kb in both sites. Taking the cytotoxicity observed during previous steps into account, the mixture of 0.5 µg of Cas9 expressing plasmid and 1 µg of plasmid containing 1 Kb homology demonstrated the optimum approach to target HBB in HEK293T cells. Doubling the length of homology did not remarkably increase the efficiency of genome editing. The PAM sequence downstream to the gRNA binding site was replaced with HindIII recognition site by mutagenized HDR template. Thus, the strategy allowed us to protect the donor HDR DNA re-cleavage of Cas9 after HDR event in the HBB gene. In addition, this strategy allows us to use insertion of HindIII into gDNA at HBB gene as a marker for HDR event at target locus. Finally, HBB long-gRNA used in these studies is effective and SCD specific, that induce high level of indels and works well in the induction of HDR based gene editing of HBB.

## 4. Discussion

Several million people in the world are affected with sickle cell disease and there is no definitive cure for it (Demirci et al., 2018). Currently, allogeneic bone marrow and hematopoietic stem cell transplantations are the only successful methods of treatment for SCD patients, yet most of the patients (>80%) don’t have a chance to find suitable donor. Moreover, even in the present of donor there is a risk for graft rejection, graft-versus-host disease and transplant related mortality. Thus, there is a need for development of novel definitive treatment approaches.

The emergence of genome-editing and gene correction technologies allowed scientists to acquire new tools in the battle against genetic diseases. These technologies have been grown up rapidly in the last 10 years. After the discovery of ZFNs and TALENs, the invention of CRISPR/Cas9 genome-editing technology has revolutionized how we approach gene editing. Thus, CRISPR/Cas9 technology is now used frequently in research labs due to the ease of design and effectiveness (Yucel and Kocabas, 2017).

The era of engineered nuclease-based genome editing has brought a new hope for inherited disorders. Using this technology, cures, not just transient and inefficient treatments, could be provided to the patients. Sickle cell disease is one of the inherited disorders, longing for a cure based on gene editing technology. Since it is caused by a point mutation on HBB gene, it is possible to efficiently induce the base change to restore functional beta subunit of hemoglobin. However, genome editing protocols still require improvements to reach 100% targeting efficiency and require lower cost, user friendly application and infinitesimal levels genotoxicity.

Here, we used CRISPR/Cas9 gene editing methodology to create DSBs in HBB locus in a human cell line. After successfully generation of DSBs in HBB locus of HEK293T cell line, we used HDR template to target and modify the locus of interest. For these purpose, two different gRNAs were designed to target the close proximity of SCD mutation. The first gRNA was a standard gRNA with 20 bp long spacer sequence with 12 bp distance to the SCD point mutation. The second gRNA named HBB long-gRNA was 25 bp in size. Long gRNA was designed by addition of extra 7 bases at the 5’ end of first gRNA to increase on-target efficiency. HBB long-gRNA also included SCD specific mutation thus it’s designed to have SCD specificity. The effect of gRNA length was investigated with the on-target mutagenesis analysis using T7E assay, and no significant difference was observed in terms of full-length and long gRNA activities on target locus. Indeed, use of full-length or longer gRNA is thought to be better for gene editing strategies when effectiveness is a key concern (Zhang et al., 2016). It was reported that longer gRNA sequences would decrease the off-target effects due to increased specificity of matching to on-target (Zhang et al., 2016).

Two different sizes of HDR templates were generated and tested in the present of SCR7 small molecules. SCR7 is small molecule and inhibitor of DNA Ligase IV, which has important role in NHEJ mechanism. Previous studies have reported that 1 µM SCR7 treatment to iPSCs resulted in significant increase in HDR mediated gene editing, inhibiting NHEJ mechanism (Chu et al., 2015). After co-delivery of gene editing components into HEK293T cells, transfection efficiencies were evaluated using flow cytometry. Transfection efficiencies were in a range between 50-60%. It was observed that increased total amount of plasmids resulted in increased cell death due to higher amounts of PEI used for transfection.

Presence of PAM sequence in the HDR template causes re-cutting by Cas9 nuclease and indel formation even after correction of desired mutation in the target location (Zhang et al., 2014). Thus, we have developed tools to remove PAM sequence as a proof-of-concept. In-fusion cloning system was utilized to remove PAM sequences in wild type HDR templates through a site directed mutagenesis. The PAM sequence downstream of gRNA binding site was replaced with HindIII recognition sequence to avoid Cas9 cleavage activity on HDR templates. Besides, HindIII sequence was used as a marker of gene editing on HEK293T cells. In case of successful gene editing via HDR mechanism, we would expect to observe cleavage on the PCR amplified target locus when treated with HindIII.

Various studies targeted HBB gene with different gene editing methodologies in various types of cell and cell lines (Antony et al., 2018; Cai et al., 2018; Chattong et al., 2017; Dever et al., 2016; Hoban et al., 2016a; Vakulskas et al., 2018; Wattanapanitch et al., 2018; Ye et al., 2016). However, our study is the first study testing long gRNA including SCD specific mutation and its effects in HBB gene editing. This study provides proof of concept that using long version of gRNAs effective in target disruption or correction in human cells. It is also noteworthy that long-gRNA with SCD mutation would likely to have less off-target activity when used in temporal manner since longer gRNA has higher affinity to and more specificity to on-target locus. In addition, the two HDR donor templates in circular form (1 Kb and 2 Kb) were generated in this study led to similar gene editing efficacy. These templates could be used to cover all possible mutations reported for HBB. On-target analysis results demonstrated the successful induction of nucleotide replacements on target locus. All combinations of Cas9/gRNA and HDR template plasmids have induced HDR based genome editing. Moreover, these findings suggest that the size of HDR templates doesn’t affect the gene editing; instead, ratio of HDR to Cas9 vector is critical in lipid based transfection methods for optimal gene editing. Further optimizations and off-target analysis are still needed to make it feasible for hard-to-transfect cells and to translate to clinical applications.

## Acknowledgments

F.K. was supported by The Marie Curie Action COFUND of the 7th Framework Programme (FP7) of the European Commission and TÜBİTAK [project no: 115C039], Turkish Hematology Association 2016 Research Project Funding, TÜBİ AK [#115S185, 215Z069, 216S317 and T 215Z071], The International Centre for Genetic Engineering and Biotechnology - Early Career Return Grant [#CRP/TUR15-02_EC], Science Academy Young Scientists award 2015 BAGEP Program, and MMV Pathogenbox Award. We like to thank Merve Uslu for technical assistance in flow cytometric analysis.

## Author Contributions

B.M.K, and F.K. designed the experiments and prepared figures. E.Y.K. contributed to gRNA cloning and transfection. B.M.K., M.Y., M.K.A. and F.K. wrote the article. All authors declare that they have no conflicts of interest concerning this work.

## Supplementary Files

**Figure S1.**
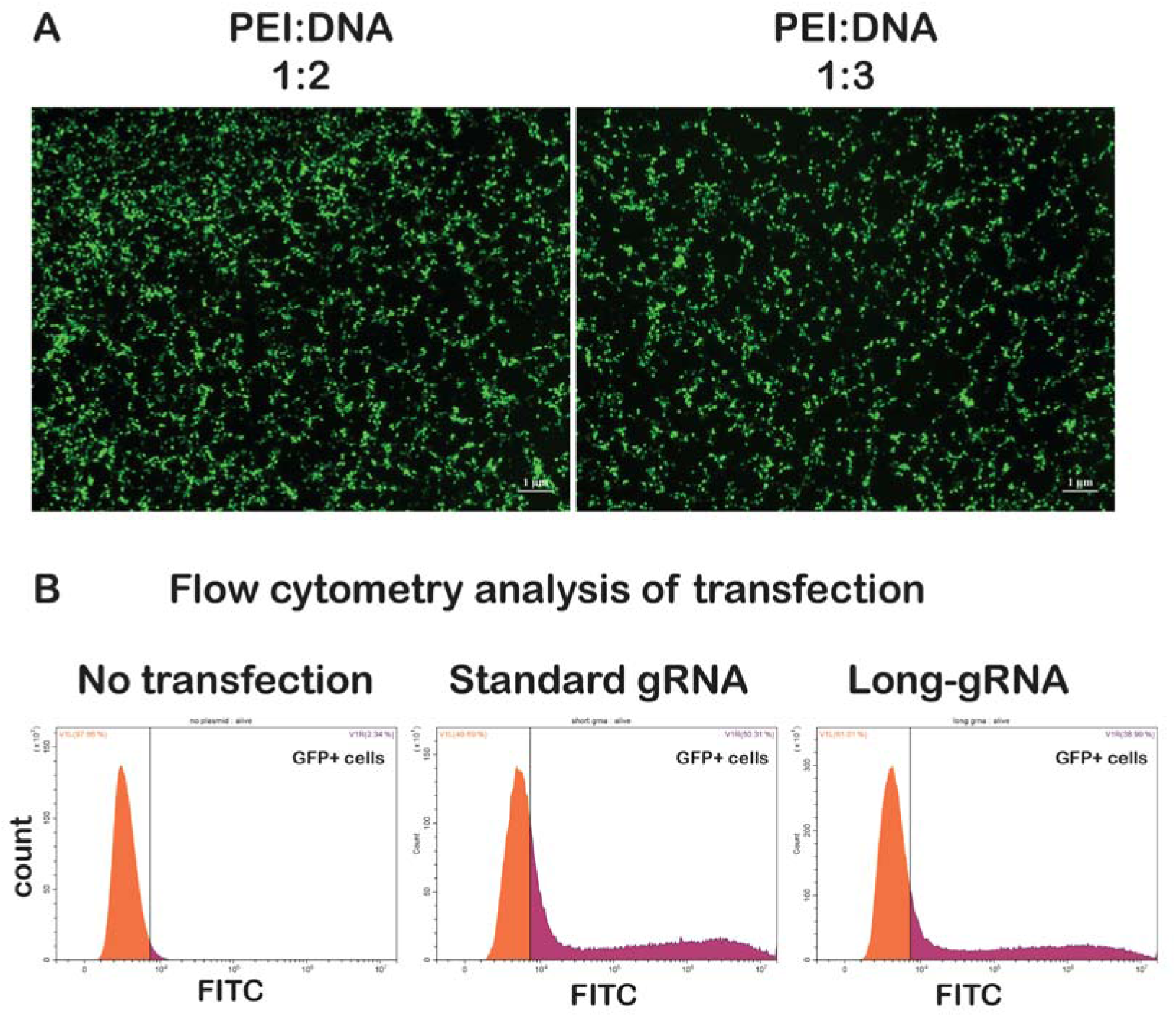
(related to Figure 2). Transfection of pSpCas9(BB)-2A-GFP Vector. **A)** Transfection of pSpCas9(BB)-2A-GFP vector into HEK293T cells were analyzed by fluorescence microscopy. PEI with two different DNA:PEI ratios were used for transfection. **B)** Flow cytometer analysis of GFP expressing HEK293T cells transfected with pSpCas9(BB)-2A-GFP plasmids transfected with PEI.

**Figure S2.**
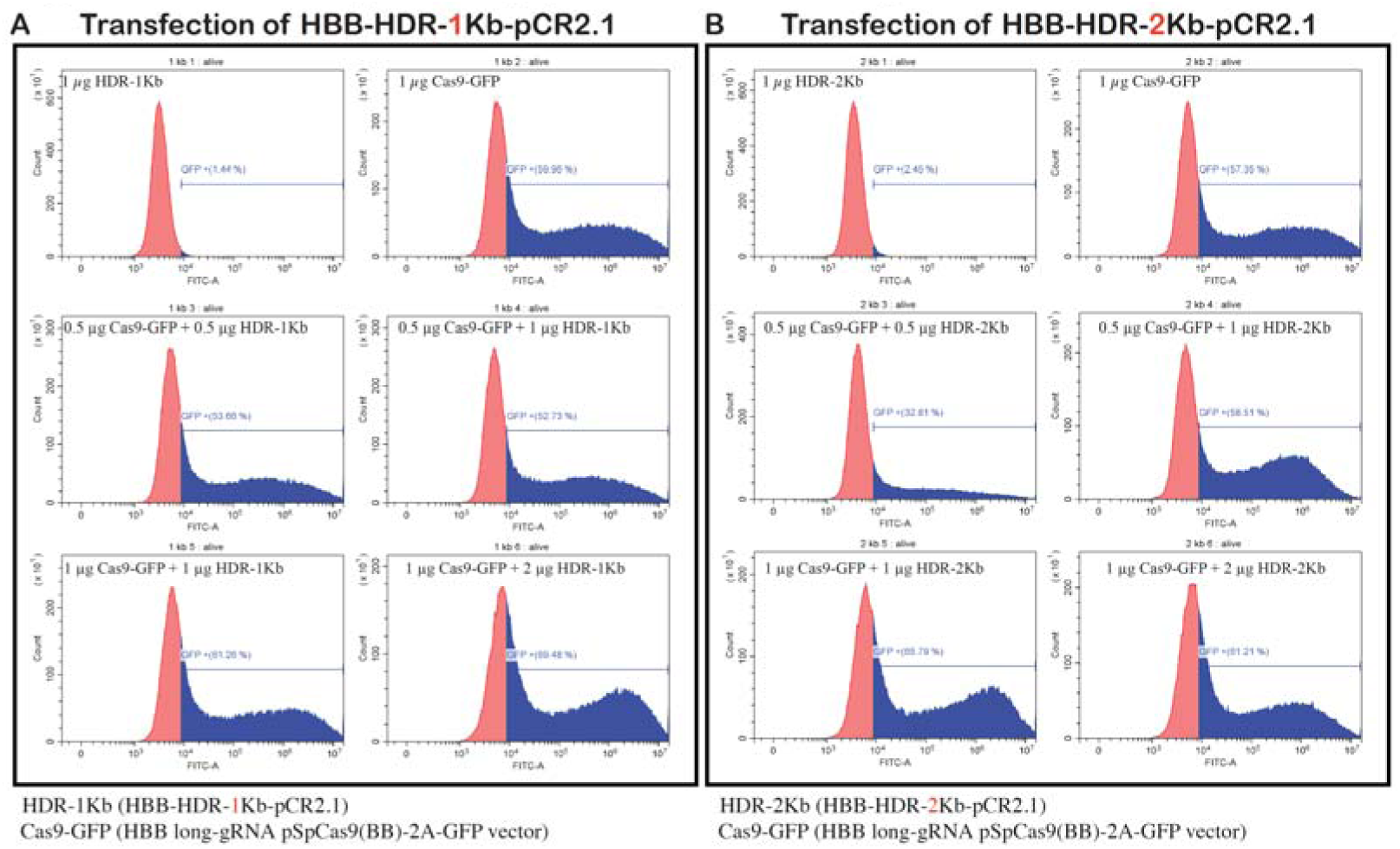
(related to Figure 5). Flow cytometry plots of HDR template transfections. Flow cytometry analysis of GFP expression post-24 hours of transfection with pSpCas9(BB)-2A-GFP +HBB-long and **A)**1 kb HDR template or **B)** 2 kb HDR template. HDR-1Kb: HBB-HDR-1Kb-pCR2.1, HDR-2Kb: HBB-HDR-2Kb-pCR2.1, and Cas9-GFP: HBB long-gRNA pSpCas9(BB)-2A-GFP vector.

**Supplementary File 1. Off-target analysis by Cas-OFFinder.** HBB standard gRNA and long gRNA were subjected to off-target prediction with Cas-OFFinder tool. Numbers of off-target were compared for every given parameter. DNA bulge, RNA bulge 0,1 and 2 were characterized along with upto 9 mismatches to determine potential targets. We have determined up to 99% reduction by HBB long gRNA compared to standard gRNA off-targets predicted

